# Scellseg: a style-aware cell instance segmentation tool with pre-training and contrastive fine-tuning

**DOI:** 10.1101/2021.12.19.473392

**Authors:** Dejin Xun, Deheng Chen, Yitian Zhou, Volker M. Lauschke, Rui Wang, Yi Wang

## Abstract

Deep learning-based cell segmentation is increasingly utilized in cell biology and molecular pathology, due to massive accumulation of diverse large-scale datasets and excellent progress in cell representation. However, the development of specialized algorithms has long been hampered by a paucity of annotated training data, whereas the performance of generalist algorithm was limited without experiment-specific calibration. Here, we present a deep learning-based tool called Scellseg consisted of novel pre-trained network architecture and contrastive fine-tuning strategy. In comparison to four commonly used algorithms, Scellseg outperformed others in average precision and Aggregated Jaccard Index on three disparate datasets. Interestingly, we found that eight images are sufficient for model tuning to achieve satisfied performance based on a shot data scale experiment. We also developed a graphical user interface integrated with functions of annotation, fine-tuning and inference, that allows biologists to easily specialize their self-adaptive segmentation model for analyzing images at the single-cell level.

## Introduction

Image-based single cell profiling is widely used in biological, pharmaceutical and medical applications, including in quantitative cytometry^1^, spatial transcriptomics^2^, high-content drug screening^3^ and cancer metastasis analysis^4^. However, due to a lack of robust and facile algorithms for single-cell analysis, average profiling remains as the most commonly used method which may cause loss of information and mislead interpretation of feature associations^5^. In recent years, deep learning has revolutionized the field of computer vision^6^ and catalyzed the advancement of single cell segmentation methods.

Differences across cell types, microscopy instruments, treatment methods, imaging modalities, and staining protocols can generate cell images with considerable diversity. As a consequence, cell segmentation algorithms were mostly developed for specific datasets^7-9^ and these methods performed poorly when applied to other styles of cell images. To overcome this limitation, generalist algorithms have been developed. In 2018, a data science bowl challenge tried to segment nuclei from a large number of different styles of microscopy images using 841 diverse images containing 37,333 nuclei^10^. Inspired by this competition, Stringer et al. annotated 608 images containing more than 70,000 segmented objects and developed a generalist algorithm named Cellpose, which exhibited excellent performance in segmenting cell bodies from many image styles^11^. Although deep-learning based generalist algorithms outperformed compared to traditional machine learning approaches like logistic regression and Random Forest (RF), the state-of-art segmentation tools still lack of capability to be self-adaptive for all kinds of cellular images. Therefore, transfer learning of segmentation models from certain source domain to in-house datasets remains an important challenge for biologists with few computational knowledges.

Fine-tuning of pre-trained models has been successfully used in computer vision^12-15^ and natural language processing^16-17^ due to its lower input requirements and more rapid convergence to a better performance. For cell instance segmentation, only few preliminary attempts were reported, such as fine-tuning of a nuclear segmentation model to satisfy different needs from distinct laboratories^18^, or transferring a pre-trained model of in vitro images to in situ tissue images^19^. However, these studies used only nuclei images for pre-training and performed binary or multiclass classification instead of instance topological maps, hence are hard to capture enough prior knowledge for fine-tuning on different kinds of cell images. Besides, they primarily tested model transferability on different nuclei images, specialized evaluation datasets for various cell-like instances such as C. elegans^20^, are far from well developed and studied. Hence, the development of a high-performance universal computational pipeline based on the fine-tuning of pre-trained models remains a challenging but important objective in automated image analysis.

In this work, we established a fine-tuning pipeline for cell segmentation algorithms and present a style-aware cell segmentation architecture named Scellseg based on attention mechanisms and hierarchical information to improve the extraction and utilization of style features. We furthermore incorporate a contrastive learning strategy to leverage information from unlabelled and pre-trained data. To evaluate the generalizability of the pipeline, we benchmarked our model on three fundamentally different styles of data from C. elegans, label-free phase-contrast cell images, and sub-cellular organelles. Furthermore, it is our first effort to estimate the minimal extent of data required for a satisfying fine-tune model and to demonstrate how instance representation and pre-trained datasets can influence model transferability. To facilitate uptake of this pipeline, we developed a graphical user interface (GUI) which can conduct annotation, fine-tuning and inference, thus making the model accessible for a wide range of users without coding experience. The model can be found at https://github.com/cellimnet/scellseg-publish.

## Results

### Design of Scellseg with pre-trained architecture and contrastive fine-tuning

Firstly, we established a pre-trained and fine-tuning pipeline for the cell segmentation model. For initial training, we utilized a dataset containing various cell types to build a generalist model. Generally, segmentation of untrained images by this model will exhibit limited performance, such as fail of detecting instances or boundary of segmentation instance. To improve the specificity of model and avoid time-consuming re-training, several images from novel data pool can be annotated for fine-tuning established model using few new labelled data (shot data) (Fig. 1a). This workflow generated a style-aware structure to better extract and comprehend style-related information and developed a new fine-tuning strategy based on contrastive learning to better make use of diverse data features, including the unlabelled data (query data) and pre-trained data. The resulting model, which we named Scellseg, contains two branches, a mask branch to compute the segmentation map of input and a contrast branch to explore the information between three types of data (Fig. 1b). The mask branch is utilized during pre-training, fine-tuning and inference, whereas the contrast branch worked only during fine-tuning.

**Fig. 1.**
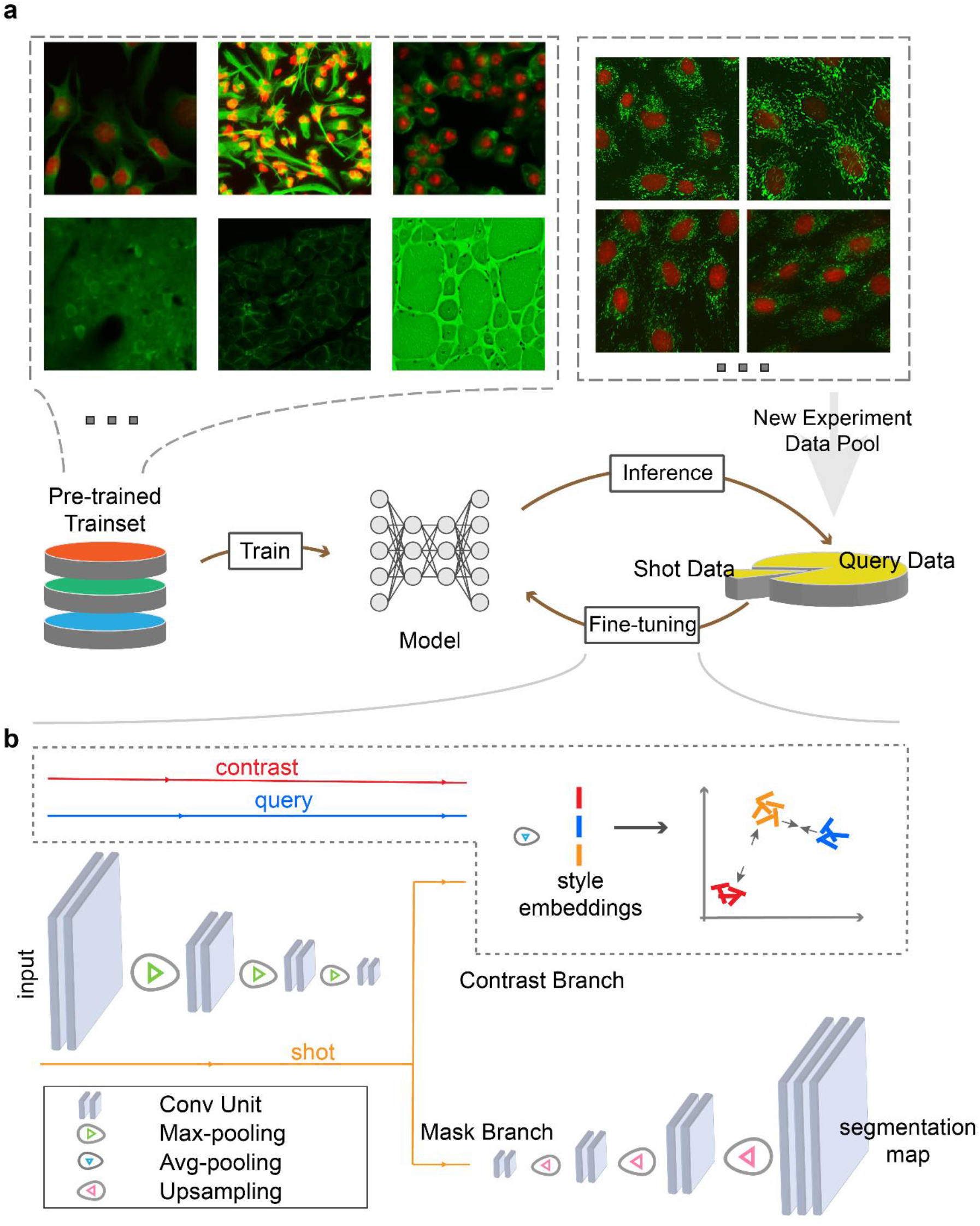
Pipeline of pre-trained and fine-tuning strategy. **a**, Overview of fine-tuning a pre-trained model for a new experiment. The shot data means hand-labelled data while query data means unlabelled data. **b**, Diagram of the proposed contrastive fine-tuning strategy. The contrast data is a subset of pre-trained data. The network is a representation of U-Net family, the detailed architecture of our proposed model is shown in Extended Data Fig. 1. Different colored lines and arrows mark the flow of data.

The mask branch was based on Cellpose model which is a member of the U-Net^21^ family of algorithms that consist of a downsampling pass that extracts features from input data, an upsampling pass that organizes different features to fit for the final task, and a concatenation operation that relays the information extracted from downsampling process to the upsampling pass. For convenient adjustment, we re-divided this structure, the last Conv Unit was split from upsampling pass and named as Tasker, and the left was named as Extractor, containing the downsampling, upsampling and concatenation parts. To improve the sensitivity of model for different styles, we added attention gates (AGs) when passing the features extracted from downsampling to the upsampling pass. These AGs give the feature map weights to highlight salient features useful for a specific task and suppress feature activation in irrelevant regions. We used dense units to consider the information from early upsampling layers, aiming to delineate accurate object boundaries. To consider different-level style information, we also fed corresponding hierarchical style embeddings into different-level dense units (Extended Data Fig. 1).

Unlike conventional fine-tuning strategies only use labelled data, to augment data utilization, we developed a contrastive fine-tuning (CFT) method to employ information from either labelled data or unlabelled and pre-trained data based on contrastive learning. Seven common cellular styles of images were chosen from pre-trained data to form contrast data (Extended Data Fig. 2). In the contrastive fine-tuning process, the contrast branch is used to compute the respective style embeddings of these three data and then a contrast loss function was designed to minimize the difference between embeddings of shot and query data from the same experiment while maximizing the difference between embeddings of shot and contrast data (Fig. 1b). This contrast loss was added into the segmentation loss function, then the total loss optimizes the model via backpropagation.

### Transferability of Scellseg with contrastive fine-tuning strategy on three evaluation datasets

To compare the performance of Scellseg in the transferability of models with other algorithms, we adopted three difference datasets named BBBC010_elegans^20^, LIVECell_bv2^22^, and mito (in-house prepared dataset containing mitochondrial images), representing three levels of cell-like images from model organism, cells to subcellular structures (Fig. 2a). In total, the datasets contained 230 images and 91,024 segmentation objects. We visualized the distribution of areas and numbers for cells per image (Extended Data Fig. 3). The average areas for three datasets are about 1000, 150 and 100,000, and numbers of instance in each image ranged from 2 to 2,815. We used t-distributed stochastic neighbor embedding^23^ (t-SNE) to visualize the style embeddings (see definition in ref.^11^) of these evaluation datasets together with pre-trained datasets and noted that the style of data in each dataset was determinant in major cluster (Fig. 2b).

**Fig. 2.**
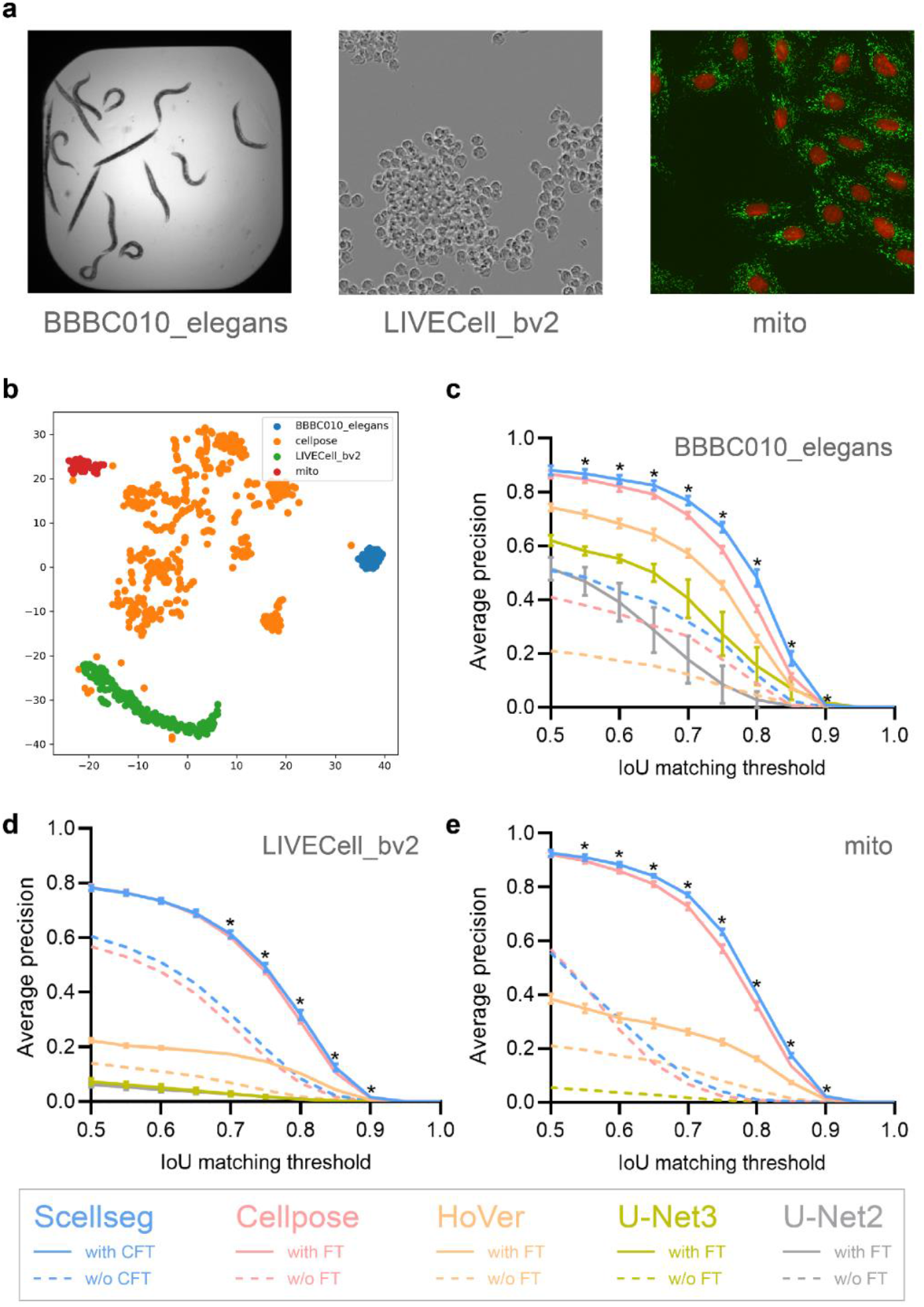
Transferability of Scellseg with contrastive fine-tuning strategy on three evaluation datasets. **a**, Example images of three datasets. **b**, Visualization for style embeddings of three datasets and pre-trained dataset using t-SNE. **c**-**e**, Performance of different models on BBBC010_elegans (**c**), LIVECell_bv2 (**d**) and mito (**e**) dataset. Different colors correspond to different models, the dotted lines denote the performance of applying models directly and the solid lines denote the performance after fine-tuning. For Scellseg, we use contrastive fine-tuning (CFT) and for others is classic fine-tuning strategy (FT). We did not plot the line which corresponding performance is less than 0.01. Each pre-trained and fine-tuning pipeline was conducted 10 times at various random states, error bars represent the mean ± SD. * means P-value<0.05, determined by two-way ANOVA followed by Sidak’s multiple comparisons test for Scellseg with CFT and Cellpose with FT.

To compare the influence of different instance representation, we benchmarked Scellseg against four other models, U-Net2^21^, U-Net3^24^, HoVer^25^ and Cellpose^11^. These four models were set with identical network structure and pre-trained with the same datasets and training strategies. We used the training data of each dataset to fine-tune the model at ten different random states, most models achieved great improvements after fine-tuning, for the BBBC010_elegans dataset, all models yielded at least 35% higher average precision. For U-Net3 model, fine-tuning strategy even yielded a dramatic increase of 62.1% in average precision. The different models differed drastically in performance, and representation of Cellpose (used in Cellpose and Scellseg model) outperformed other methods. Scellseg with contrastive fine-tuning achieved the best performance on all three datasets, especially on the BBBC010_elegans dataset. At the universally used intersection over union (IoU) threshold of 0.5, our Scellseg and Cellpose both achieved high average precision when segmenting C. elegans (0.882 for Scellseg; 0.868 for Cellpose), microglial cell BV-2 (0.783 for Scellseg; 0.784 for Cellpose), and mitochondria in cardiomyocytes (0.927 for Scellseg; 0.922 for Cellpose). In contrast, Scellseg performed considerably better at higher thresholds, such as 0.75 on all three data sets ([0.670, 0.493, 0.634] for Scellseg compared to [0.587, 0.475, 0.571] for Cellpose, respectively; Fig. 2c-e).

We also compared the performance of Scellseg with or without contrastive fine-tuning strategy. As shown in Fig. 3, it is clear that the fine-tuned model exhibited considerably better capability of instance detection. Importantly, fine-tuning strategy improved the ability of distinguishing adjacent cells, which allowed the segmentation of scattered mitochondria around the nuclei in mito dataset. Furthermore, our contrastive fine-tuning strategy outperformed the classic method on the Cellpose test set after fine-tuning on the three evaluation datasets. However, as expected, all re-trained models suffered a sharp decline compared with the initial generalization ability (Extended Data Fig. 4).

**Fig. 3.**
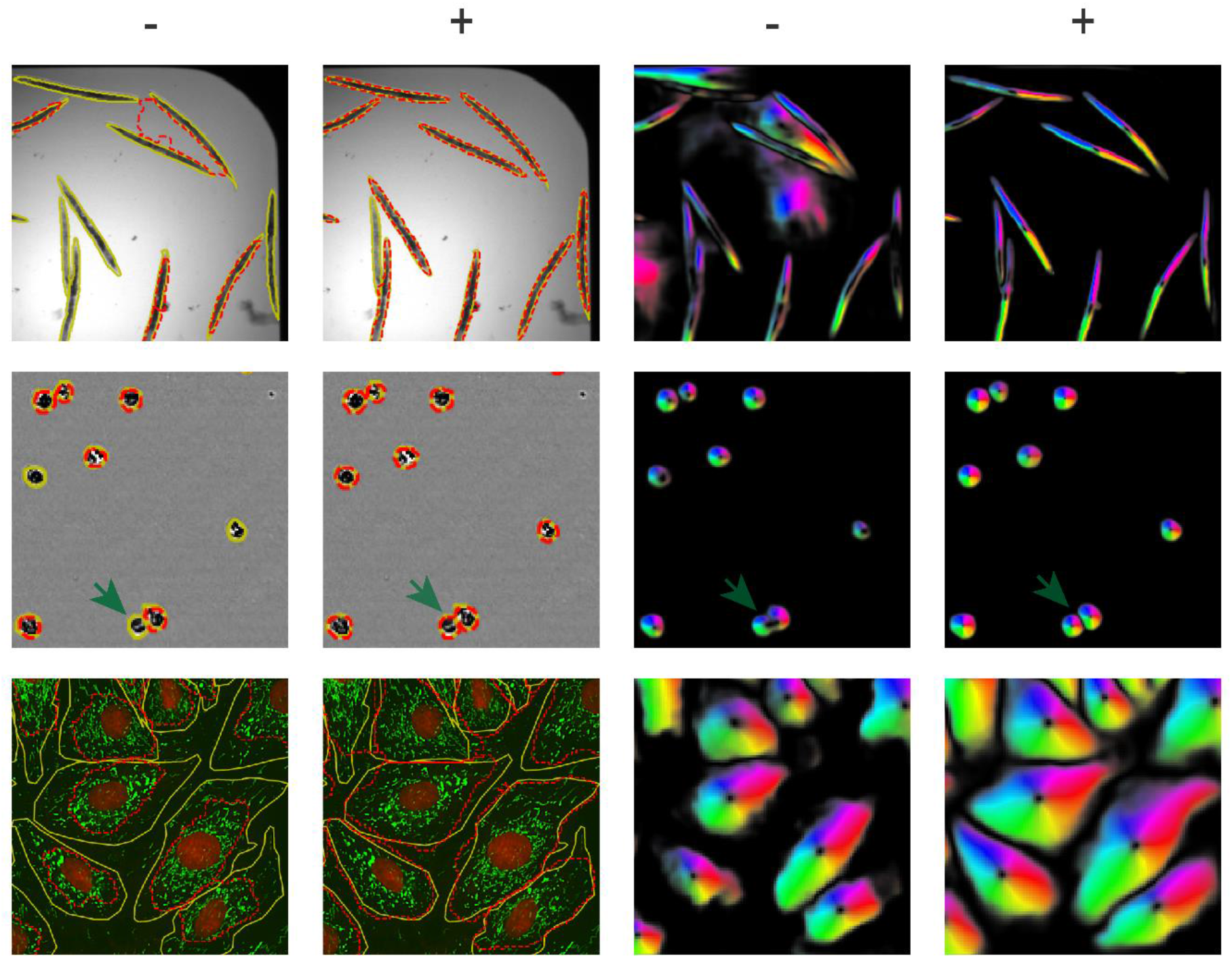
Example Scellseg segmentation results with and without contrastive fine-tuning on three evaluation datasets. The right two columns show the direct topological maps outputted by Scellseg model. The left two columns show ultimate masks of different datasets, the ground truth masks are shown in yellow solid line, and the predicted masks are shown in dotted red line. Symbol “−” represents results of applying models directly and symbol “+” represents results after contrastive fine-tuning. Green arrows emphasize the segmentation of adherent cells.

### Pre-trained dataset scale experiments

To explore how the pre-trained dataset can influence model transferability, we used different subsets of the Cellpose training set. The initial subset (Sneuro) only contains one style of images from the Cell Image Library^26^, then additional styles of images were sequentially added, such as fluorescent cells (Sfluor), non-fluorescent and membrane-labelled cells (Scell), other microscopy data (Smicro), as well as non-microscopy images (“Sgeneral”, corresponding to the full Cellpose train set; Fig. 4a).

**Fig. 4.**
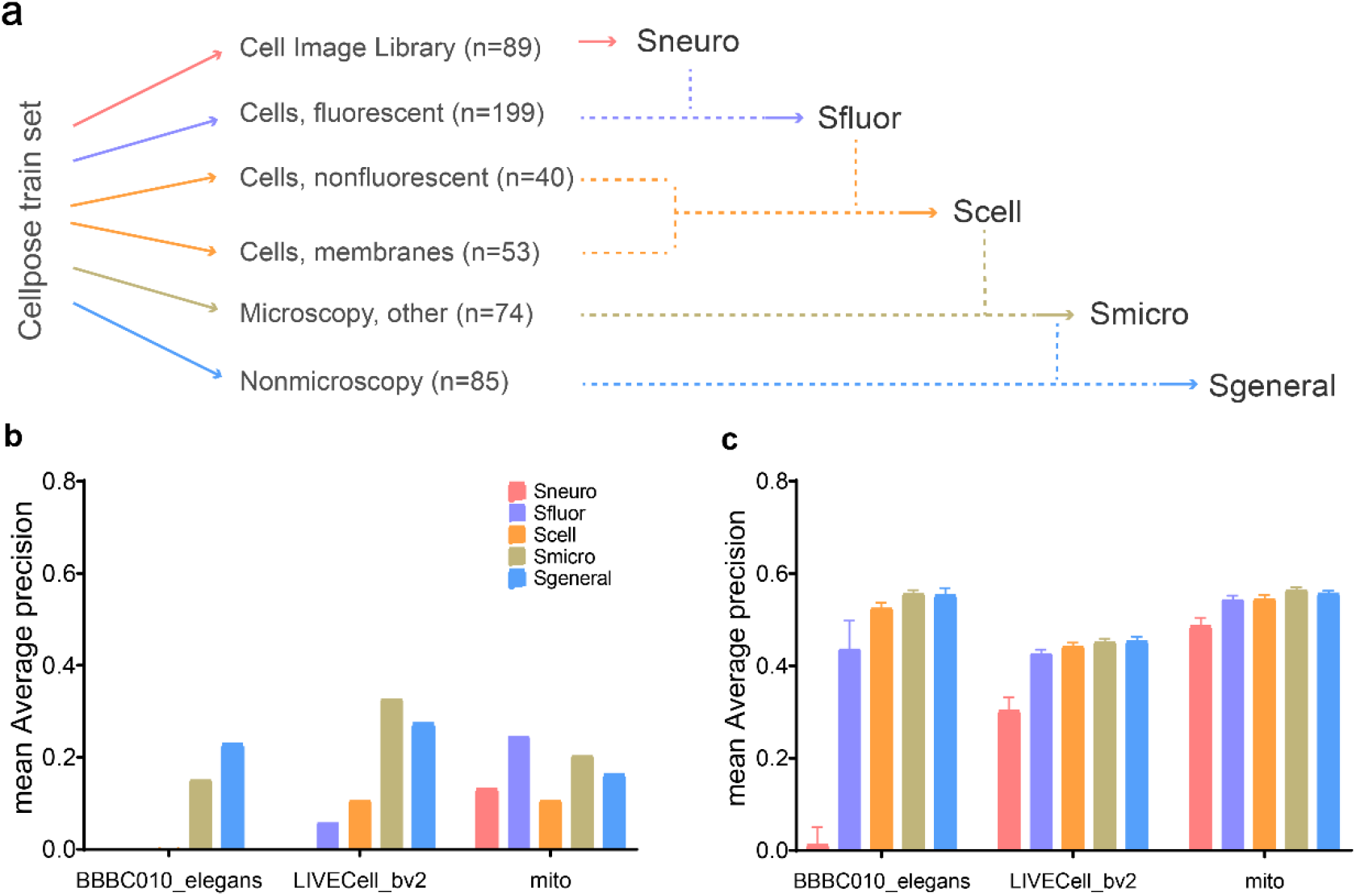
Pre-trained dataset scale experiments. **a**, Composition of different subsets from Cellpose train set. **b**, Generalization ability of different pre-trained Scellseg models on three evaluation datasets. Generalization ability means the performance of employing pre-trained model directly. **c**, Transferability of different pre-trained Scellseg models on three evaluation datasets. Transferability means the performance of employing the pre-trained model after fine-tuning. Each pre-trained and fine-tuning pipeline was conducted 10 times at various random states, error bars represent the mean ± SD.

We pre-trained Scellseg with Sneuro, Sfluor, Scell, Smicro and firstly tested the generalization ability of each model by applying it directly without any adaptation on three evaluation datasets. For C. elegans, the model trained with Sneuro, Sfluor, and Scell does not result in successful recognition until the pre-trained dataset contains microscopy instances with structures beyond cells. For small and round BV-2 cells, the generalization ability also increased with the richness of the dataset from Sneuro to Smicro, and model trained on Smicro even outperformed the model trained on Sgeneral. For the segmentation of mitochondria, surprisingly the model trained with Sfluor and Smicro outperformed all others (Fig. 4b).

Next, we tested the transferability of each model (Fig. 4c). As expected, the transferability of Scellseg increased with the richness of the dataset from Sneuro to Smicro, and Scellseg pre-trained with Smicro achieved similar transfer performance on three evaluation datasets compared to Scellseg pre-trained with Sgeneral ([0.555, 0.451, 0.564] and [0.554, 0.454, 0.557], mean average precision [mAP] of Smicro and Sgeneral respectively). On the BBBC010_elegans dataset, a model pre-trained on Sneuro achieved only very poor transfer performance (0.013 mAP) and performance increased substantially only after the addition of different styles of fluorescent images (0.436 mAP). As more cell-like images added, performance increased further to 0.554 with the full pre-trained dataset.

### Shot data scale experiments and ablation experiments

To explore the extent of annotated data required for fine-tuning, we made a shot data scale experiment on these evaluation datasets. We set 10 scale levels, and for each shot number, we randomly sampled 10 times from the training pool to fine-tune the model, followed by testing of transferability. For this evaluation, we focused on Scellseg with CFT and Cellpose with classical fine-tuning because these models clearly outperformed the other three algorithms. For all datasets, we observed that initial performance is relatively low with large variance (Fig. 5a). As the number of shot images increases, the performance improves drastically and variance becomes smaller while Scellseg significantly outperformed Cellpose. For BBBC010_elegans, LIVECell_bv2 and mito dataset, Scellseg with CFT get [2.0%, 4.8%, 18.5%] final improvement respectively compared to [4%, 6%, 12.2%] for Cellpose with classical fine-tuning. We conducted curve fitting using Hyperbola function for each method per dataset for further inspection of the transferability across different shot numbers. The results show that, when increasing shot number, different methods converged to different values and the mAP converged differently across datasets. For mito dataset, mAP persistently increased while for BBBC010_elegans, the rate of its convergence is relatively fast, and whatever the dataset is, performance plateaued at eight shots. Similar results were obtained using the mean Aggregated Jaccard Index^27^ as a means to evaluate transfer performance (Extended Data Fig. 5). Therefore, it is suggested that at least of eight images is required to achieve satisfied transfer learning based on generalized model.

**Fig. 5.**
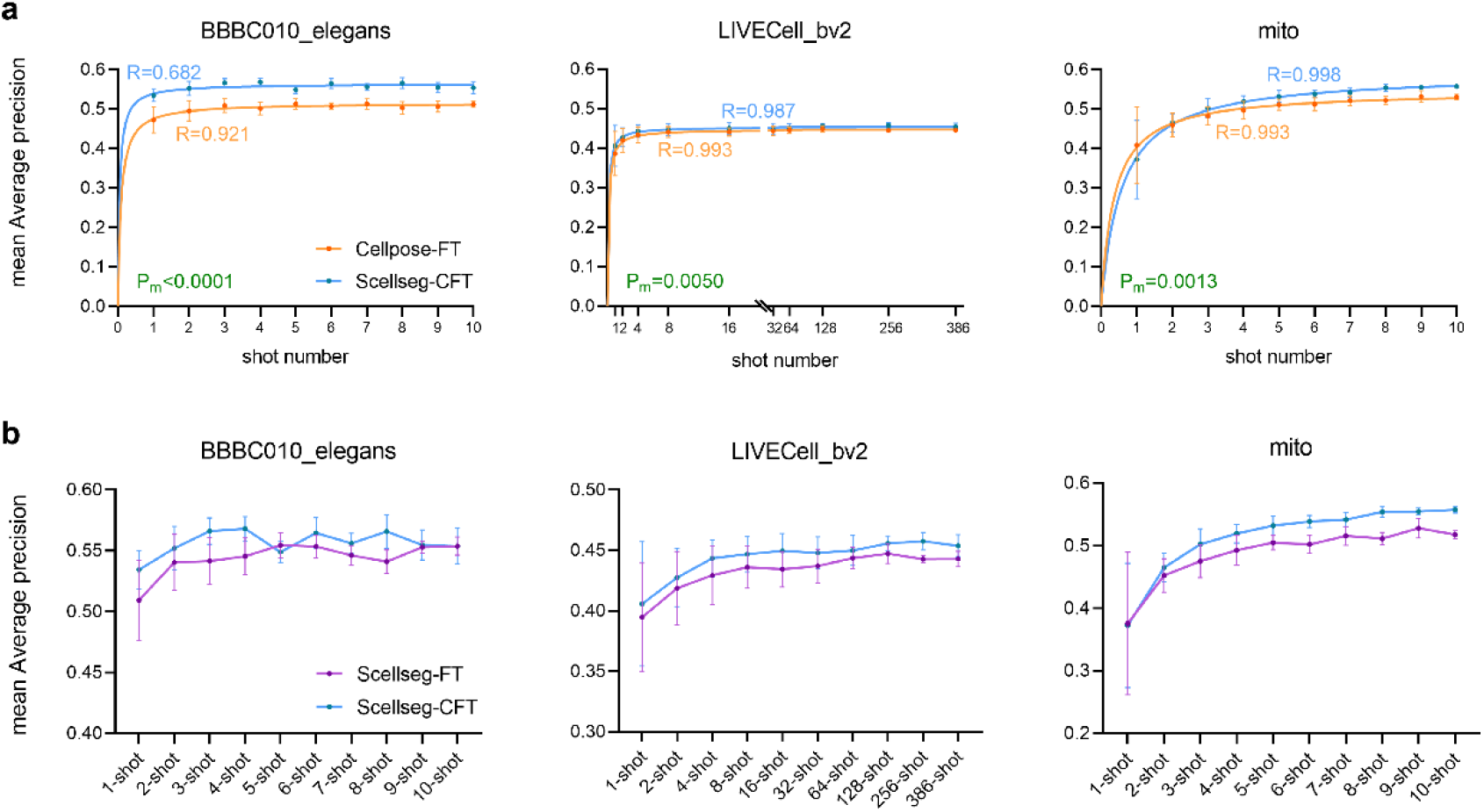
Shot data scale experiments (a) and ablation experiments (b). Each pre-trained and fine-tuning pipeline was conducted 10 times at various random states, error bars represent the mean ± SD. **a**, Performance of Cellpose-FT and Scellseg-CFT on three evaluation datasets at 10 shot data scales. Cellpose-FT represents Cellpose with classic fine-tuning and Scellseg-CFT represents Scellseg with contrastive fine-tuning strategy. Another metric to evaluate the models is shown in Extended Data Fig. 5. We performed nonlinear regression based on hyperbola function and corresponding R-value of fitted curve is plotted in the picture. A two-way ANOVA analysis was conducted for group comparison of Scellseg-CFT and Cellpose-FT per dataset and corresponding Pm-value is plotted in the picture. Pm-value<0.05 was considered the performance between Scellseg-CFT and Cellpose-FT is significant. **b**, Ablation experiments for contrastive fine-tuning strategy. Scellseg-FT represents Scellseg with classic fine-tuning strategy, data of Scellseg-CFT is completely same as Scellseg-CFT in (**a**).

To verify the function of our contrastive fine-tuning strategy, we conducted ablation experiments. Importantly, our contrastive fine-tuning strategy outperformed Scellseg using the classic fine-tuning method at different shot-number experiments on all three evaluation datasets (Fig. 5b). Due to the similar performance of the model when “only” trained on Smicro, we conducted the same shot data scale experiments and ablation experiments as Scellseg pre-trained on the Sgeneral dataset and, excitingly, our style-aware pipeline worked and again outperformed Cellpose with classic fine-tuning strategy (Extended Data Fig. 6).

### Graphical user interface

To facilitate Scellseg accessible for scientists without coding experience, we designed a GUI (Fig. 6) with three functional parts, i) view and draw, ii) fine-tune, and iii) inference. For basic annotations, users can modify the mask of an instance directly at single-pixel resolution without deleting the whole mask. We also developed a cell list management system to help users edit the corresponding mask and provide annotations, thereby allowing to provide ground truth for segmentation and cell class prediction simultaneously. Furthermore, users can easily save or load cell lists.

**Fig. 6.**
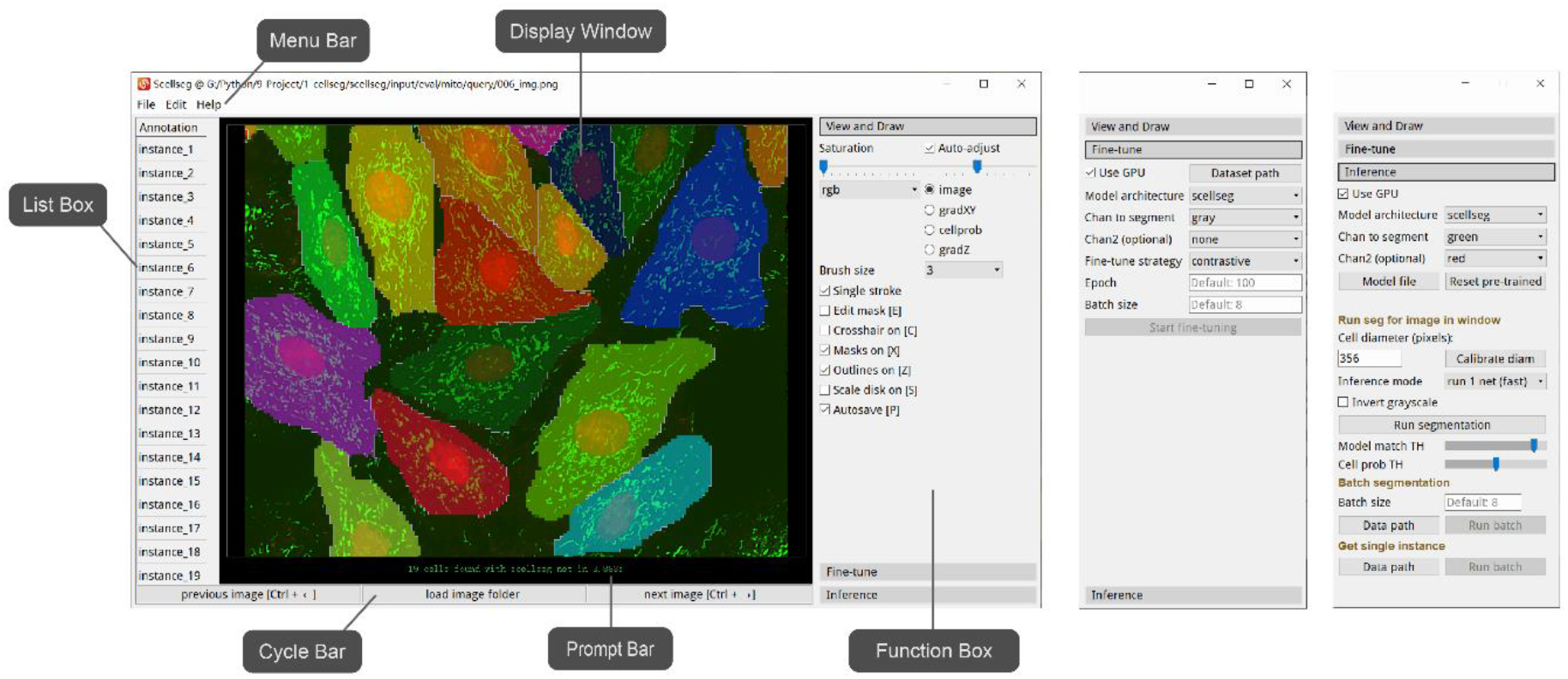
Graphical user interface (GUI). This GUI contains six modules: Menu Bar, List Box, Display Window, Prompt Bar, Cycle Bar and Function Box. There are three main functions: “View and draw”, “Fine-tune” and “Inference”.

In the second module, users can fine-tune the pre-trained model with their own manually labelled data. The system allows users to choose a pre-trained model from Scellseg, Cellpose and HoVer. Furthermore, each model can be combined with either the contrastive or classic fine-tuning strategy, presented above. It will not only give biologists and pathologists more flexibility and versatility for their image analysis tasks, but also help algorithm engineers to easily conduct experiments to study such pre-trained and fine-tuning pipelines. Finally, users can use the fine-tuned model to conduct inference either for one image or use batch inference. After annotation or inference, users can also acquire images of each single instance for further analysis.

## Discussion

Accurate cell instance segmentation is still a challenging task for many laboratories. Although generalization models have been developed, these typically require large annotated datasets, which is time- and labor-consuming in data collection, particularly when a large number of segmented objectives are supposed to be covered. To augment data utilization, we firstly established a pipeline for the fine-tuning of pre-trained cell segmentation algorithms. On this basis, we proposed a style-aware pipeline, yielding the best transferability on three different benchmarking datasets. Specifically, the work achieved three main innovations: Firstly, we refined the architecture of Cellpose through introducing attention mechanisms and hierarchical information, making the model more sensitive to different styles. Secondly, we implemented a contrastive fine-tuning strategy to leverage the information from both unlabelled and pre-trained data based on contrastive learning, which have also achieved great success in other deep learning applications^28-32^. Finally, we organized three benchmarking datasets containing three levels of cell images for further use in segmentation algorithm development.

Importantly, after fine-tuning, AP for BBBC010_elegans and mito can reach up to about 0.9 at a threshold of 0.5, more than 37% and 36% improvements respectively, resulting in model performance that is generally acceptable for researchers to conduct reliable downstream analysis. Our benchmarking showed that among four different instance representations, topological maps generated by the Cellpose model constituted the best way to introduce rich instance information. Results could be further improved by our style-aware pipeline, which exhibited the best transferability on all three evaluation datasets, indicating that introducing such style relevant information can benefit the fine-tuning process. Notably, while generalization ability overall declined after fine-tuning to a specific task, our contrastive fine-tuning considerably improved generalizability. Furthermore, emerging methods like continual learning^33^ are also worth to investigate in this context.

In the pre-trained dataset scale experiments, we observed that transferability of Scellseg-CFT increased with the richness of the pre-trained dataset, suggesting that our Scellseg-CFT pipeline can also benefit from large-scale and high-diversity datasets. Notably, the generalization ability of Scellseg pre-trained with Sfluor containing different styles of fluorescent images, outperformed all other models on the mito evaluation dataset, indicating that model pre-training on diverse but specialized data may yield greater performance than both low-diversity specialized dataset (such as Sneuro) or high-diversity generalized datasets (such as Scell that also contains nonfluorescent images). However, we did not observe similar phenomena on transferability. We also noted that the generalization ability increased with the addition of more different microscopy instances beyond cells to other non-cell instances like C. elegans, again demonstrating the success of our style-aware pipeline.

For the shot data scale experiment, it is not surprising that performance increases along with shot number. However, what excited us is that we observed a large payoff when increasing shot number from 1 to 3, whereas performance plateaued after approximately eight shots. These results are of high practical relevance as they indicate that the annotation of only about eight images is sufficient to yield a sufficiently fine-tuned model. Few-shot^34^, one-shot^35^ and zero-shot^36^ learning strategies can be studied to further reduce the number of annotated images needed. Notably, at small shot numbers, different shot data can have very large impacts on the fine-tuning process, whereas we observed that as the shot number increases variance becomes substantially smaller. In the future, active learning^37^ on cell instance segmentation promises to refine shot data selection for fine-tuning.

In this work, we did not research the influence of basic model backbone and all models were based on convolutional neural networks (CNN). In recent years, self-attention architectures (such as Transformer^38^) have shown great success and there have been studies attempting to apply them to computer vision^39^. Such transformer architectures have better expressive ability but require more data for accurate training. Nevertheless, we believe that such approaches will eventually provide an important improvement in computer vision compared to CNN.

By integrating attention mechanisms and hierarchical information for style-aware segmentation with a contrastive fine-tuning strategy, Scellseg features the highest transferability when benchmarked on three diverse imaging datasets against currently used segmentation methods. Scellseg optimizes cell and object recognition in diverse microscopy data and, combined with an easy-to-use GUI, can make advanced parallelized segmentation accessible also to researchers and histologists without coding experience. Moreover, the Extractor and Tasker design can facilitate the adaption to other computer vision tasks, such as segmentation and simultaneous class prediction^25^, or conducting feature extraction for phenomics analysis^40^. We anticipate Scellseg will serve not only for cell segmentation, but also other for a wide range of other applications in cell biology and biomedicine.

## Methods

The code was written in Python programming language v.3.7.4. All experiments were conducted on NVIDIA GeForce RTX 2080Ti. The deep learning framework used Pytorch^41^ v.1.7.1.

### Datasets

#### Pre-training datasets

We used the Cellpose dataset published by Stringer et al^11^ which contains a total of 608 images and over 70,000 segmented instances. 540 images were used as training set (the last of every 8 images was chosen as validation set) and 68 images were used as test set. Here, the whole training set (also named as Sgeneral) was used to pre-train the models and the test set was used to evaluate the generalization ability in Extended Data Fig. 4. Furthermore, a subset of the training set containing a total of seven styles of images with five images per style was used as the contrast data (Extended Data Fig. 2).

#### Evaluation datasets

Three datasets were used to evaluate the transferability of different models, here called BBBC010_elegans, LIVECell_bv2 and mito. BBBC010_elegans was downloaded from the Broad Bioimage Benchmark Collection^42^, containing 100 images of *C. elegans* in a screen to find novel anti-infectives. There are two phenotypes in this dataset, for worms treated with ampicillin, they appear curved in shape and smooth in texture, while untreated worms appear rod-like in shape and slightly uneven in texture. Only the brightfield channel was used. We discarded images with heavily crossed instances because it is not the focus of our work, the problem may be solved by some special postprocessing algorithm^20^ or introducing the z-axis information when designing the ground truth. Finally, 49 images were reserved, 10 were used as the training set and 39 were used for testing.

The LIVECell_bv2 dataset^22^ consists of 536 phase-contrast images and over 330,000 segmented instances. These images were achieved using label-free phase-contrast imaging and cells in this dataset have small spherical morphology and are homogeneous across populations. Of the available images, 386 were used as the training set and 152 were used for testing.

We also generated a novel dataset called Mito dataset, which consisted of 49 fluorescent images of mitochondria from high content screening studies. The images were acquired by ImageXpress Micro Confocal (Molecular Devices). Each image contains two distinct channels, a nuclear channel stained with Hoechst-33342 (Sigma) and a mitochondria channel stained with tetramethylrhodamine methyl ester (TMRM, Sigma). All these images were manually annotated by a single human operator (D.J.X.), 10 images were used as training set and 39 were reserved for testing. Because there was no clear boundary between individual cells, the Mito dataset was used to compare the performance of different algorithms regarding mitochondrial segmentation at the single cell level.

All these three datasets were organized in Cellpose format. The summary information can be seen in Extended Data Table. 1.

### Models

When training a cell instance segmentation model, we usually provide raw images and the corresponding masks which label the individual instances with different positive integers per image. Although different values can represent different instances in these masks, it is impractical to directly predict such masks because the max value in each mask represents the number of instances, which are different across images. Thus, the model has to pre-set a very large shape of last conv unit in Pytorch^41^ tensor shape format to cover all instances. However, such approaches can result in inefficient memory usage and may not learn well in such a high dimensionality. It is challenging to find an excellent representation of instances and until now there have been four main methods: U-Net2^21^, U-Net3^24^, HoVer^25^ and Cellpose^11^.

For the classic U-Net model (usually called as U-Net2), we directly map the annotated masks to 2-classes, zero represents background and one represents instance. This method usually performs poorly on touching cells because instance information was completely discarded. In 2018, Fidel et al^24^ introduced cell borders as the third class to make the network notice the original gap between cells (usually called as U-Net3), they yield a significant improvement compared with U-Net2. In 2019, Simon et al^25^ further developed the model on multiple independent multi-tissue histology image datasets. For each cell per image, they generated horizontal and vertical distance maps to bring in rich instance information when inference, marker-controlled watershed^43^ was used as the postprocessing to create the final masks. In 2020, Stringer et al^11^ generated topological maps through a process of simulated diffusion from masks, and when on a test image, they used gradient tracking^44^ to recover individual cells.

Here, we wanted to compare how expressive power different methods can provide, so we used the same architecture as Cellpose, only changed the final shape of convolutional layer, loss function and postprocessing to adapt to each method. Scellseg model adopted the representation of Cellpose because the best performance in the experiments. Expect the different parts of Scellseg, all other architecture sets were same as Cellpose too.

Two-channel 224×224 images were set as input for all 5 models in this work. The primary channel contains instances to segment and the second optional channel can provide extra information such as nuclei channel to support model learning. The hierarchical level of Conv Units was set as [32, 64, 128, 256]. We computed the style embeddings through applying the average pooling on feature map of last Conv Unit and the dimensionality of each level style embeddings after being concatenated in upsampling pass is [256, 384, 448, 480].

### Pre-train segmentation models

#### Pre-train different models with Sgeneral

We trained five models (U-Net2, U-Net3, HoVer, Cellpose, Scellseg) with Sgeneral, which contains totally 540 images, 64 of which were reserved for validation.

For U-Net2 and U-Net3, a learning rate of 0.002 was selected to achieve good model convergence. For HoVer and Scellseg, the loss function is same as Cellpose, which was defined as:

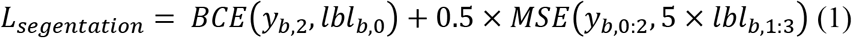

Where BCE represents the binary cross-entropy loss, MSE represents the mean square error loss, y represents the ultimate output of model, lbl represents the ground truth, and subscripts corresponded to respective dimensions in y or lbl, b represents the batch size, which we here set to 8.

All models were trained for 500 iterations with stochastic gradient descent, the mean diameter was set to 30, all other training hyper-parameters were same as Cellpose.

#### Pre-training Scellseg for pre-trained dataset scale experiments

We trained four other models across different subsets of Cellpose mentioned above: Sneuro, Sfluor, Scell, Smicro. For each subset, the last of every 8 images was reserved for validations (11, 36, 47, 56, respectively). All other training hyper-parameters were the same as for Scellseg pre-trained with Sgeneral.

Training logs of all models are shown in Extended Data Fig. 7.

### Fine-tune segmentation models

#### Classic fine-tuning strategy (FT)

When fine-tuning, batch size was set to 8, epoch was set to 100, the optimizer was Adam, the initial learning rate was set to 0.001 and every quarter of epochs it was reduced by 50%. Before being fed to the network, the image-mask pairs were resized, randomly rotated and reshaped with the ultimate shape of input as (8, 2, 224, 224).

#### Contrastive fine-tuning strategy (CFT)

The contrast loss function was defined as:

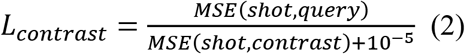

Where MSE represents the mean square error and was used to compute the difference between embeddings, and 10^-5^ was added to prevent divisions by zero. This contrast loss was added into the segmentation loss function during contrastive fine-tuning, so the final loss function was defined as:

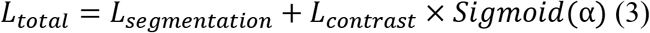

Where α is a scalar to control the weight of contrast loss, also learnt during the fine-tuning process. A sigmoid function was used to assure that the coefficient of contrast loss changed smoothly between zero and one. The initial α in the contrast loss function was set to 0.2, initial learning rate of α was set to 0.1, with reductions by 50% every quarter of epochs. For query and contrast data, they were resized, randomly cropped, randomly rotated and reshaped before being fed to the network with identical input shapes. Other parameters were the same as for the classic fine-tuning strategy.

For both classic and contrastive fine-tuning strategies, we fine-tuned all layers because this method performed best compared with downsampling part or the whole extractor (Extended Data Fig. 8). For each dataset, we computed the instance diameter using shot data without using the automated method provided by Cellpose, which was used in resizing the current mean diameter of instances to the mean diameter used for model pre-training.

### Benchmarking

#### Metrics

We used average precision (AP) and the Aggregated Jaccard index (AJI) to evaluate segmentation performance (See ref. ^11^ for detailed definitions). Except in Fig. 2c-e, we averaged the AP or AJI over IoU from 0.50 to 0.95 with a step size of 0.05 for convenient comparison and reserving the overall performance information simultaneously as detailed below:

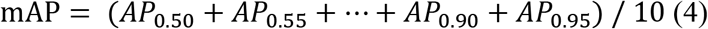

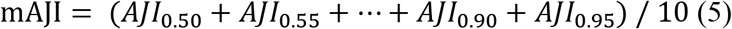

#### Shot data scale experiments

We set a total of 10 scale levels; for BBBC010_elegans and mito we used [1, 2, 3, 4, 5, 6, 7, 8, 9, 10] and for LIVECell_bv2 [1, 2, 4, 8, 16, 32, 64, 128, 256, 386]. For each shot number experiment, we randomly sampled 10 times from the training set to fine-tune the pre-trained model. To eliminate issues due to different training data, the random state was kept identical across models. For example, we sampled the 9th, 5th, and 2nd image from the total of 10 images in the training set of a 3-shot experiment for the mito dataset, and then used the same images as training data for all five models.

### Statistical Analysis

All figures were made using GraphPad PRISM 8.0 software (GraphPad Software, Inc., CA, USA). All graphs display mean values, and the error bars represent the standard deviation (SD). Statistical analyses were conducted with two-way repeated measures analysis of variance (ANOVA) followed by Sidak's multiple comparisons test in Fig. 2c-e and two-way ANOVA in Fig. 5a. A nonlinear regression curve fit was performed using a hyperbolic function in Fig. 5a, given as:

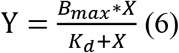

Where B_max_ and K_d_ are constants.

## Data availability

Three evaluation datasets used in this work will be made available at https://scellseg-data.s3.cn-northwest-1.amazonaws.com.cn/evaluation_datasets.zip upon publication.

## Code availability

The Scellseg software, including detailed tutorial, can be freely available at GitHub (https://github.com/cellimnet/scellseg-publish).

## Acknowledgments

The authors are grateful for the support from ZJU PII-Molecular Devices Joint Laboratory and support from "Medicine + X" interdisciplinary Center of Zhejiang University.

## Author contributions

Y.W. and R.W. proposed the concept and supervised the overall project. D.J.X. established the pipeline, organized datasets, written the code of pipeline, conducted experiments, performed data analysis, designed and finished the code writing of graphical user interface. D.H.C. participated in part of the code writing of graphical user interface. D.J.X., V.M.L., Y.W., R.W., Y.T.Z. participated in the preparation of the manuscript.

## Extended Data

**Extended Data Fig. 1.**
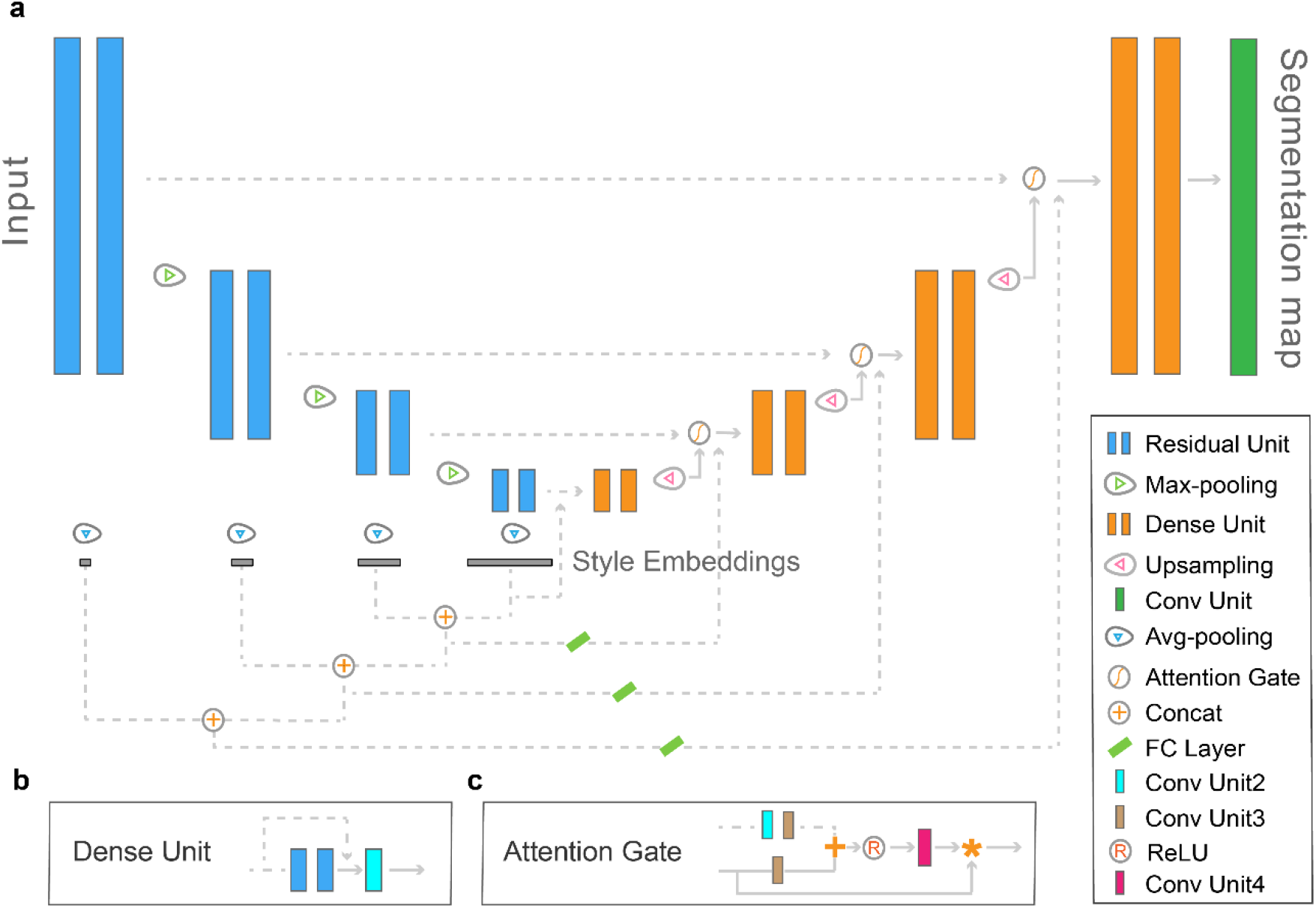
Architecture of the proposed Scellseg. **a**, The Residual Unit refers to Cellpose^11^, Dense Unit refers to Hover-Net^25^, Attention Gate refers to Attention U-Net^45^, Conv Unit represents a set of operations including BatchNorm2d, ReLU and Conv2d in Pytorch^41^. The blue parts (including Max-pooling operations) were called as downsampling pass, the dot lines (including operations marked on them) were called as concatenation part, specially, the orange parts (including Avg-pooling operations) were called as upsampling pass, and the last green Conv Unit was named as Tasker, downsampling, upsampling together with concatenation part were named as Extractor. Input images are progressively encoded and decoded to get the ultimate segmentation map. Style embeddings of each scale is obtained by using global average pooling on respective convolutional map. **b**, Detail architecture of Dense Unit. Conv Unit2 represents a set of operations including Conv2d, BatchNorm2d and ReLU. **c**, Detail architecture of Attention Gate. Conv Unit3 represents a set of operations including Conv2d and BatchNorm2d. Symbol “+” represents tensor plus and Symbol “*” represents tensor multiplication. Conv Unit4 represents a set of operations including Conv2d, BatchNorm2d and Sigmoid.

**Extended Data Fig. 2.**
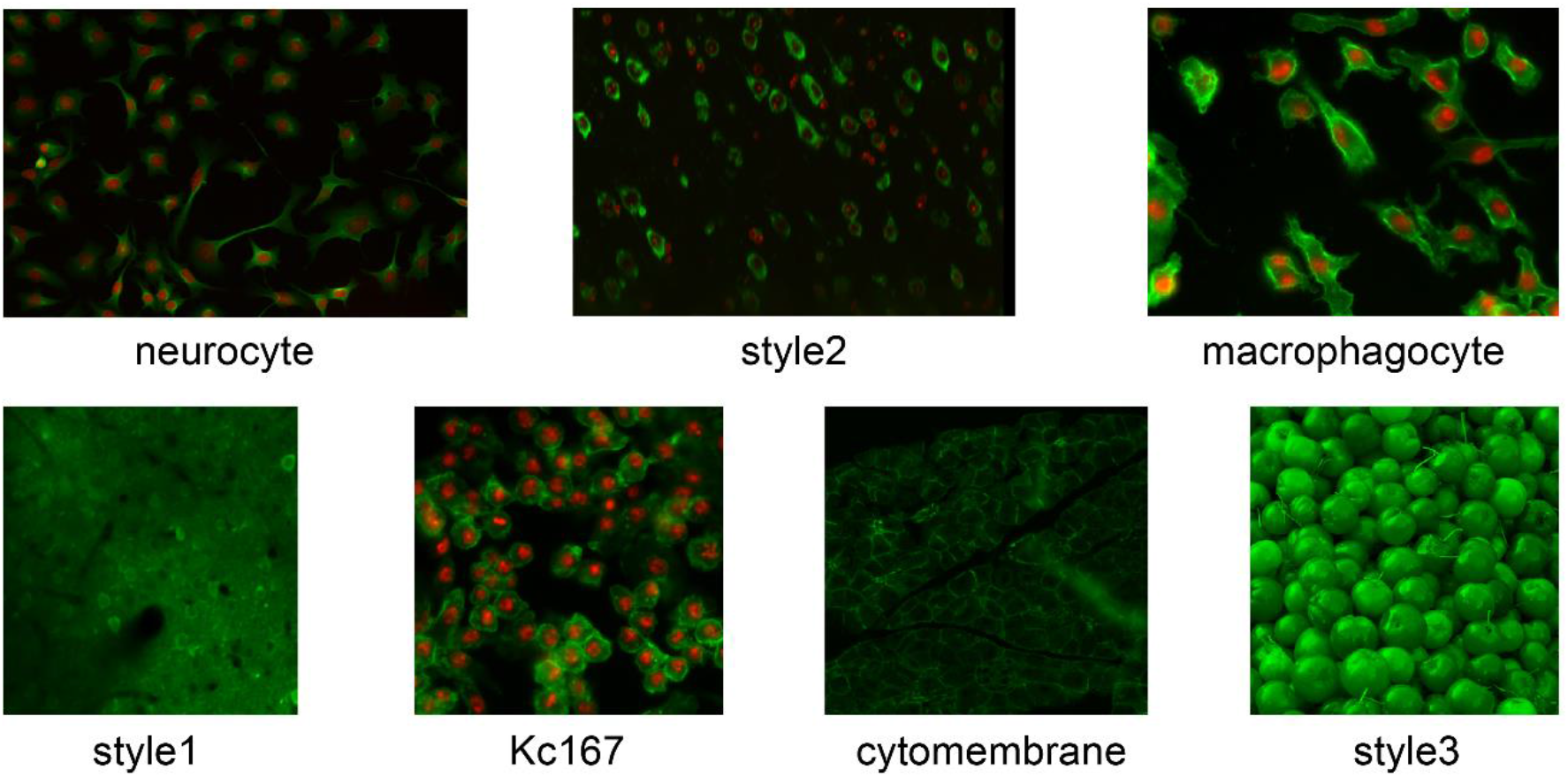
Example image per style of Contrast data. There are totally 7 styles of images we used in our contrastive fine-tuning strategy, each style includes five images.

**Extended Data Fig. 3.**
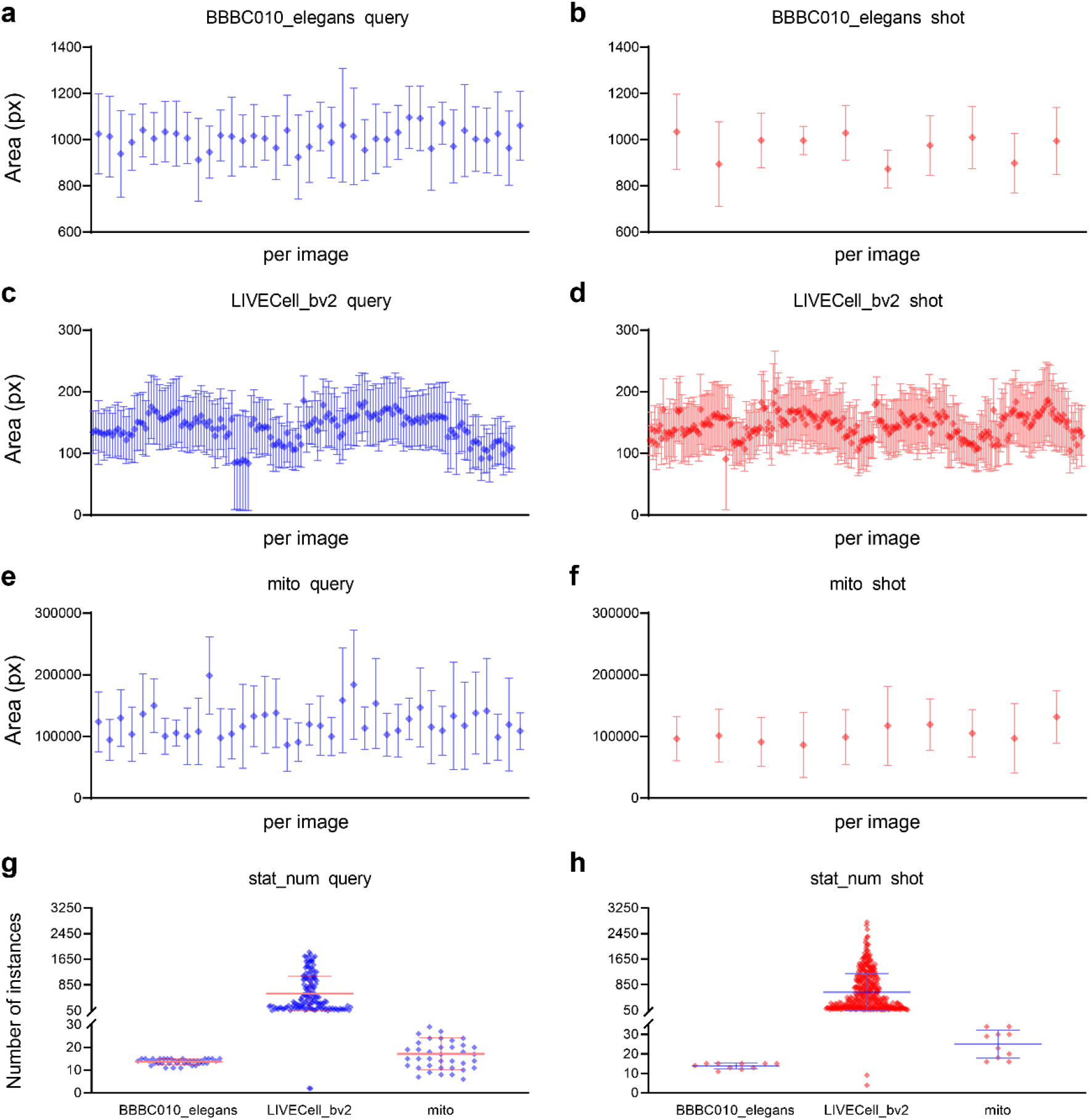
Statistics of three evaluation datasets. **a-f**, Distribution of cell areas in px for each image in query or shot data of BBBC010_elegans (**a, b**), LIVECell_bv2 (**c, d**), mito (**e, f**) dataset, error bars represent the mean ± SD. **g-h**, Distribution of number of instances for each image in query data (**g**) or shot data (**h**) of three datasets, each dot represents one image in corresponding dataset, error bars represent the mean ± SD.

**Extended Data Fig. 4.**
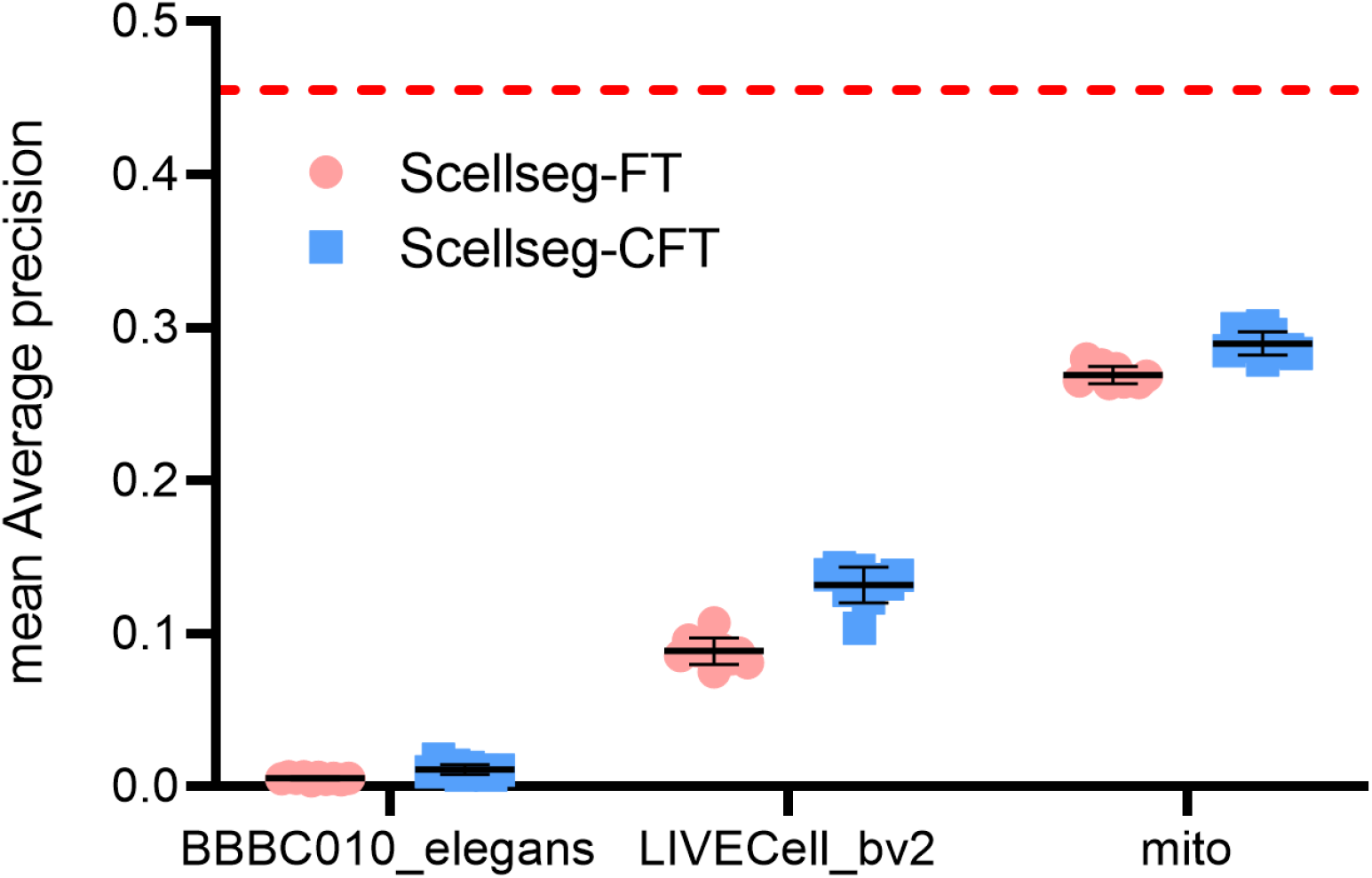
Generalization ability on Cellpose test set of Scellseg with different fine-tuning strategies. Scellseg-FT represents Scellseg with classic fine-tuning and Scellseg-CFT represents Scellseg with contrastive fine-tuning strategy. Red dot line represents employing Scellseg directly on test set. Each pre-trained and fine-tuning pipeline was conducted 10 times at various random states, error bars represent the mean ± SD.

**Extended Data Fig. 5.**
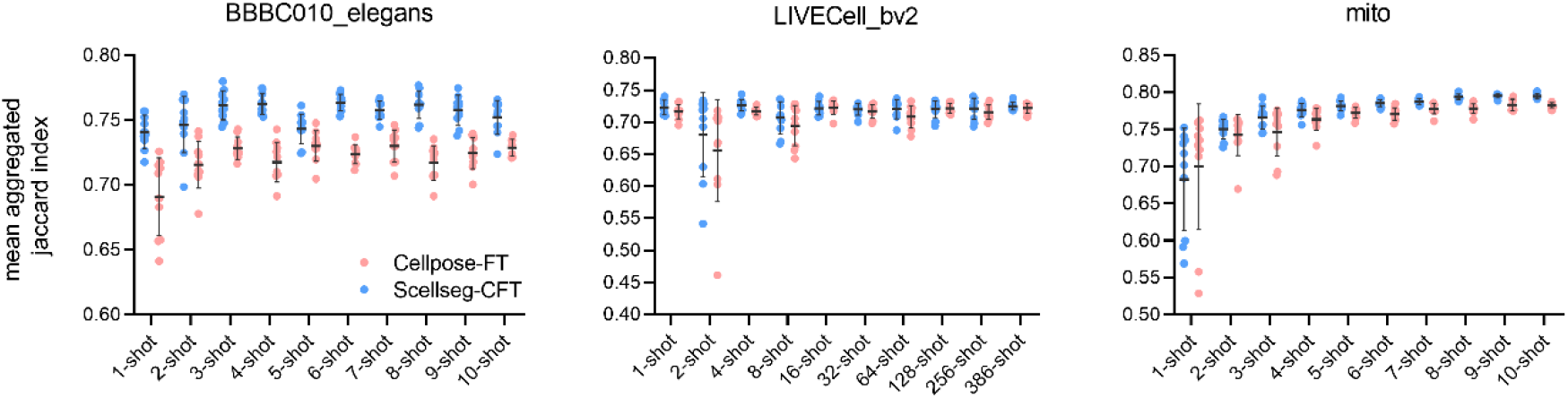
Using mean Aggregated Jaccard Index metric to evaluate segmentation performance in shot data scale experiments. Error bars represent the mean ± SD.

**Extended Data Fig. 6.**
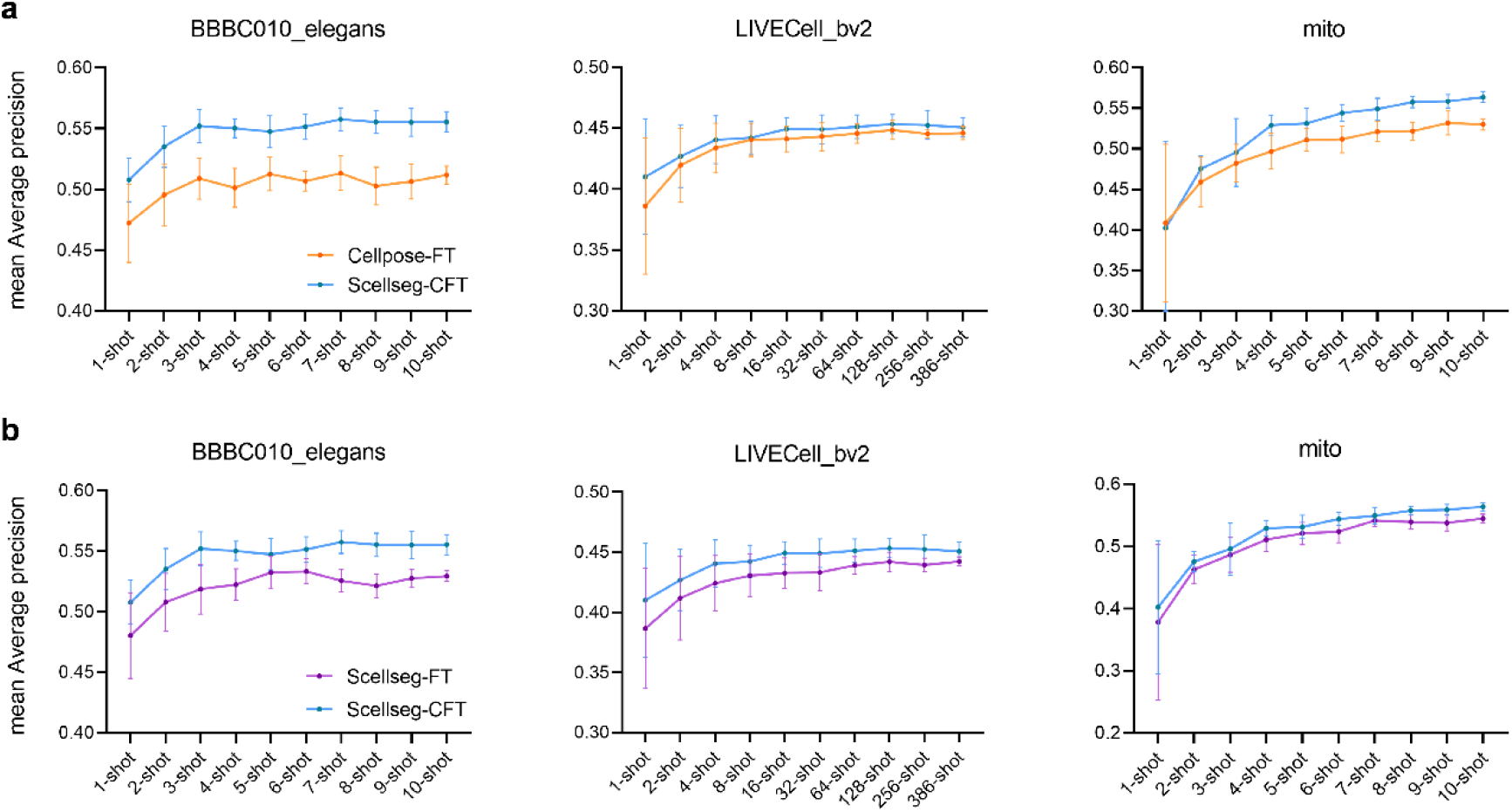
Shot data scale experiments (a) and ablation experiments (b) on Scellseg pre-trained with Smicro. Each pre-trained and fine-tuning pipeline was conducted 10 times at various random states, error bars represent the mean ± SD. Data of Scellseg-CFT in (**a**) is completely same as Scellseg-CFT in (**b**).

**Extended Data Fig. 7.**
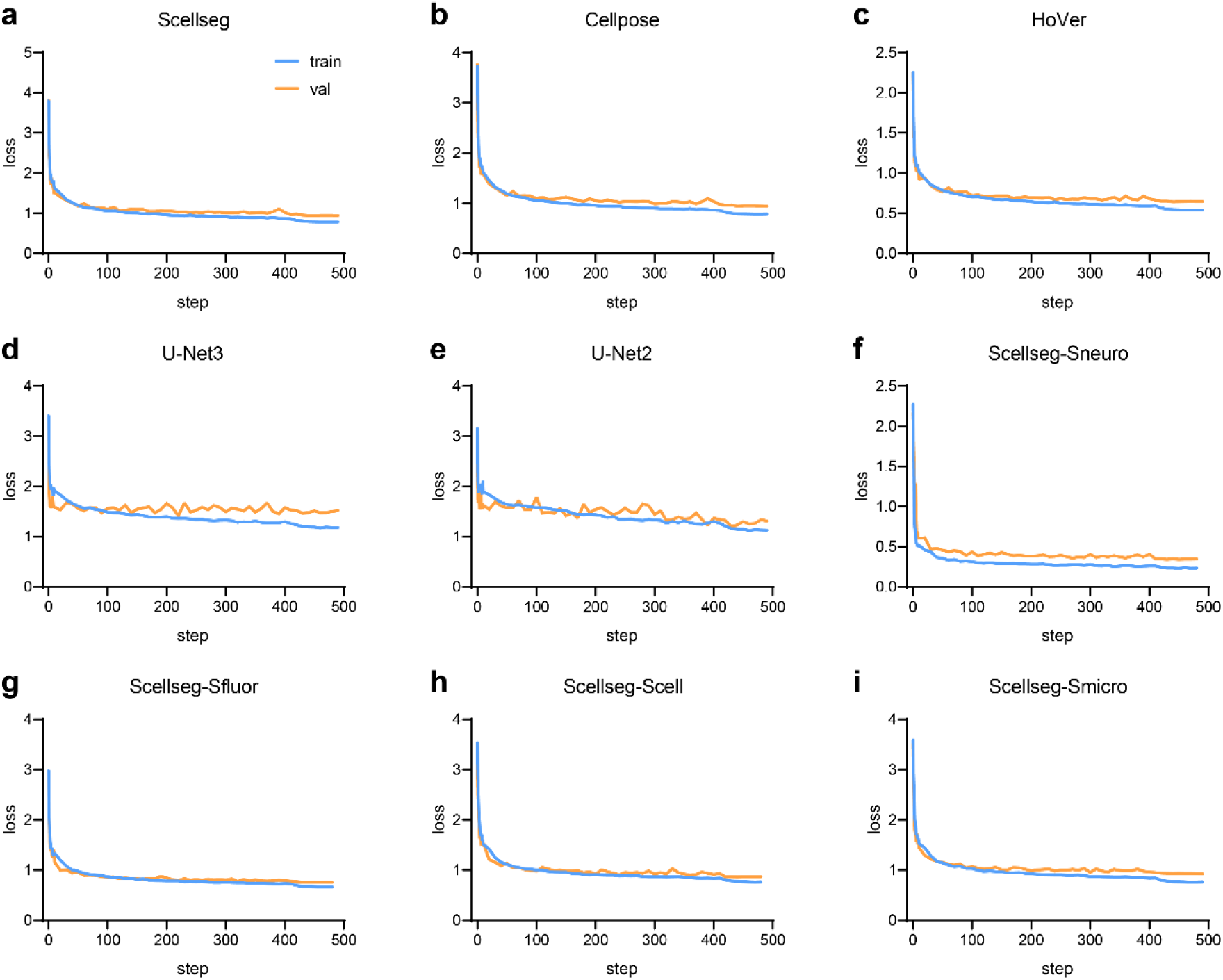
Train logs of models in the paper.

**Extended Data Fig. 8.**
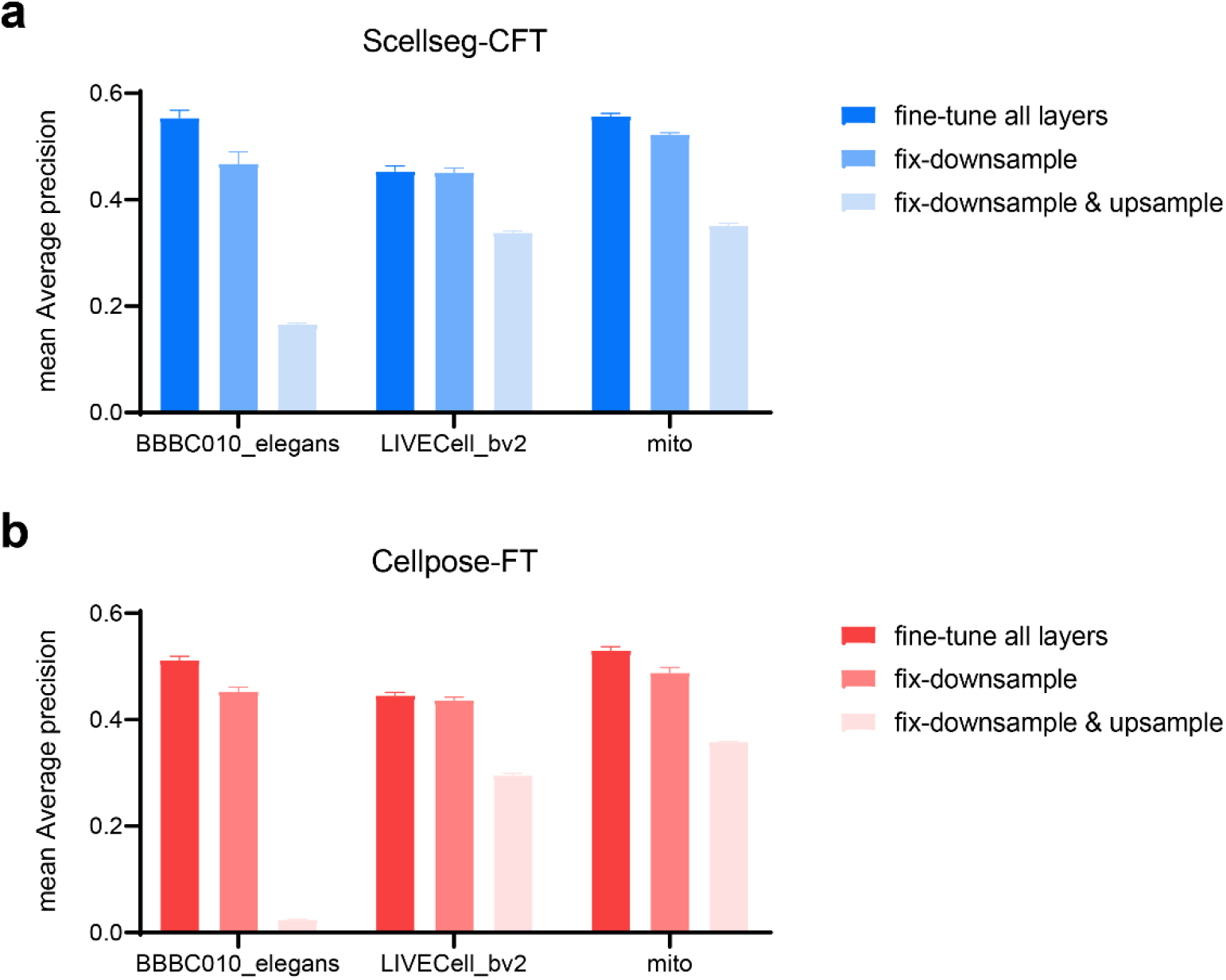
Performance of different fine-tuning methods. We compared three kinds of fine-tuning methods for Scellseg-CFT (**a**) and Cellpose-FT (**b**) on three evaluation datasets, respectively are fine-tuning all layers of the model, fixing the downsampling layers and fixing downsampling-upsampling layers. Each fine-tuning method was conducted 10 times at various random states, error bars represent the mean ± SD.

**Extended Data Table. 1.** Summary statistics of three evaluation datasets.

| Dataset | Number of images in train set | Number of images in test set | Number of instances in test set | Acquisition Modality | Shape of images | Characteristics |
| --- | --- | --- | --- | --- | --- | --- |
| BBBC010_elegans | 10 | 39 | 670 | Bright Field | 696*520*1 | worms appear rod-like or curved in shape |
| LIVECell_bv2 | 386 | 152 | 533 | Fluorescent, 10X | 704*520*1 | small spherical morphology, homogeneous population |
| mito | 10 | 39 | 89821 | Phase-contrast imaging, 60X | 2048*2048*3 | mitochondria scatter around the nuclei, no clear boundaries |
| **Total** | **406** | **230** | **91024** |  |  |  |

